# Cortactin interacts with αDystrobrevin-1 and regulates neuromuscular junction morphology

**DOI:** 10.1101/2023.10.13.562179

**Authors:** Teresa De Cicco, Marcin Pęziński, Olga Wójcicka, Klemens Rottner, Tomasz J. Prószyński

**Affiliations:** Łukasiewicz Research Network ‒ PORT Polish Center for Technology Development, Stabłowicka 147, 54-066 Wrocław, Poland; Nencki Institute of Experimental Biology, Polish Academy of Sciences, 3 Pasteur Street, 02-093 Warsaw, Poland; Division of Molecular Cell Biology, Zoological Institute, Technische Universität Braunschweig, Germany; Department of Cell Biology, Helmholtz Centre for Infection Research, Inhoffenstraße 7, 38124 Braunschweig, Germany

**Keywords:** neuromuscular junction, Dystrobrevin, dystrophin-glycoprotein complex, actin, cortactin

## Abstract

**Background:** Neuromuscular junctions allow for transmitting signals from the nervous system to skeletal muscles, triggering their contraction, and their proper organization is essential for breathing and voluntary movements. αDystrobrevin-1 is a cytoplasmic component of the dystrophin-glycoprotein complex and has pivotal functions in regulating the integrity of muscle fibres and neuromuscular junctions. Previous studies identified that αDystrobrevin-1 functions in the organization of the neuromuscular junction and that its phosphorylation in the C-terminus is required in this process.

**Methods:** We used synthetic peptides corresponding to the phosphorylated tyrosine Y730 at the C-terminal part of αDystrobrevin-1 to precipitate interacting proteins from homogenate of differentiated muscle cells. Isolated proteins were identified by mass spectrometry, and co-immunoprecipitation and bimolecular fluorescence complementation experiments in skeletal muscles were used to validate interactions. We used immunohistochemical analysis and muscle electroporation to study cortactin localization in skeletal muscles. To study the function of cortactin in the neuromuscular system, we used cortactin KO mice. Synaptic morphology was studied using unbiased automatic image analysis, and muscle strength was assessed in grip-strength experiments and an animal’s ability to run on voluntary wheels and a treadmill.

**Results:** Our proteomic screen identified a number of putative αDystrobrevin-1 interactors recruited to the Y730 site in both its phosphorylated and unphosphorylated state. Amongst various actin regulators, we identified the Arp2/3 complex regulator cortactin. We showed that similarly to αDystrobrevin-1, cortactin is strongly enriched at the neuromuscular postsynaptic machinery and obtained results suggesting that these two proteins interact in cell homogenates and at the neuromuscular junctions. Analysis of synaptic morphology cortactin knockout mice showed abnormalities in the slow-twitching soleus muscle and not in the fast-twitching tibialis. However, muscle strength examination did not reveal apparent deficits in knockout animals.

**Conclusions:** Our studies identified cortactin as a novel interactor of the dystrophin-glycoprotein complex, pivotal in maintaining muscle and neuromuscular junction integrity. We have shown that cortactin is a novel postsynaptic machinery component that can be essential in organizing the neuromuscular junctions.

## 1. Background

The Neuromuscular junction (NMJ) is a specialized synapse in the peripheral nervous system that connects axons of the motor neurons with skeletal muscle fibers. Its primary role is to allow for the transmission of signals from the nervous system to skeletal muscles, triggering muscle contraction. Proper functioning of the neuromuscular transmission controls vital functions such as voluntary movements and respiration. Therefore, malfunctions of the neuromuscular system are associated with severe disorders, including muscular dystrophies and myasthenia gravis.

In vertebrates, motor neurons release the neurotransmitter acetylcholine (ACh), which is detected by acetylcholine receptors (AChRs) at the postsynaptic site. AChRs must be clustered at a high density underneath presynaptic neurotransmitter release sites for efficient neuromuscular transmission. Complex processes, including local expression of postsynaptic components, clustering of AChR at the cell surface, and stabilization of synaptic sites, ensure enrichment and maintenance of AChR at the junctions. Local synthesis within the large muscle fibers is controlled by signals that originate from the nerve. A major player in this process is the secreted glycoprotein agrin, detected on the postsynaptic side by LDL receptor-related protein 4 (Lrp4) and Muscle-specific kinase (MuSK) [1, 2]. Interactions with agrin activate MuSK and initiate signalling that controls the expression of many postsynaptic proteins from specialized synaptic nuclei anchored at the postsynaptic machinery [1]. MuSK activation also leads to a cascade of events that stimulates the clustering of AChR upon their exocytosis [1]. Rapsyn is a primary protein responsible for aggregating AChR molecules into bigger assemblies [3]. At the same time, rapsyn interacts with various components of the cytoskeleton, contributing to the stabilization of AChR and the entire postsynaptic machinery [4, 5].

The dystrophin-associated glycoprotein complex (DGC) is a large macromolecular and multifunctional membrane-associated complex present in the sarcolemma throughout the muscle fiber. The DGC spans the sarcolemma and constitutes a crucial link between the intracellular cytoskeleton and the extracellular matrix (ECM) [6, 7]. These properties allow the DGC to also stabilize postsynaptic components and serve as scaffold for the recruitment of signalling molecules and other peripheral proteins. The DGC consists of two classes of components: the core components and the satellite proteins. The core components, including dystrophin, utrophin, dystroglycans, sarcoglycans, sarcospan, syntrophin, and dystrobrevins, form the structural foundation of the DGC [7]. On the other hand, molecules such as nNOS (neuronal nitric oxide synthase), GRB2 (growth factor receptor-bound protein 2), α-catulin, and liprin-α1 are classified as satellite peripheral components [8]. Dysfunctions of the DGC have received significant attention in recent decades due to their recognition as primary cause of a group of inherited skeletal myopathies. Among muscular dystrophies, Duchenne muscular dystrophy is the most prevalent form, and loss-of-function mutations in the dystrophin gene is the underlying cause [9].

αDystrobrevins (αDB) is a group of proteins with a partial amino acid sequence similar to dystrophin that are cytoplasmic components of the DGC in skeletal muscles. Among the existing splicing variants of αDB, αDB1 and αDB2 represent the main isoforms in vertebrates [7, 10]. αDB1 is highly concentrated at the NMJ, and αDB2 is expressed along the entire muscle fiber [7, 10, 11]. The two αDB isoforms have virtually identical amino acid sequences; however, αDB1 has an additional C-terminal sequence missing in αDB2 that contains three phosphorylated tyrosine residues [11, 12]. These tyrosine residues are essential for the synaptic functions of αDB1, which include regulation of AChR stability and NMJ morphology [12, 13]. In the absence of functional αDB1, the amount of AChR at the junctions and their stability is reduced [12, 14]. This is associated with morphological changes of NMJs, displaying ragged edges with spicules containing AChR protruding outside the junctions. However, the molecular mechanisms downstream of αDB1 phosphorylation are poorly understood. Our previous work identified several proteins that are recruited to the phosphorylated tyrosine Y713 in the C-terminal part of αDB1, and that are involved in the regulation of AChRs [15].

Here, we report the identification of proteins that interact with αDB1 tyrosine Y730. One of them is an actin cytoskeleton regulator known as cortactin (Cttn). We demonstrate that Cttn is enriched at the postsynaptic domain of the NMJ, where it colocalizes with AChR and interacts with αDB1. Using Cttn knockout mice, we show that Cttn plays an important role in the organization of the neuromuscular junction.

## 2. Materials and Methods

### 2.1. Plasmids

The EGFP-cortactin plasmid was a gift from Anna Huttenlocher (Addgene plasmid #26722; http://n2t.net/addgene:26722; RRID: Addgene_26722 [16]); pBiFC-VC155 and pBiFC-VN173 were gifts from Chang-Deng Hu (Addgene plasmid # 22011; http://n2t.net/addgene:22011; RRID:Addgene_22011 and Addgene plasmid # 22010; http://n2t.net/addgene:22010; RRID:Addgene_22010, respectively) [17]; pBiFC-αDB-VN173 was a kind gift from Mohammed Akaaboune [18]. αDB1-VN173, VC155-Cttn, and Cttn-VC155 were generated by NEBuilder® HiFi DNA Assembly Cloning Kit (# E5520S, New England BioLabs), which enables the assembly of multiple inserts within a linearized vector in a one-step PCR reaction. EGFP-cortactin and pBiFC-VC155 were used as PCR templates to obtain the fragments encoding Cttn and VC155 (155-238), and inserted into the pBiFC-VN173 backbone. Two versions of Cttn plasmid were generated, which carried VC155 fused to either N- or C-terminus of cortactin. The αDB1VN173 plasmid was generated by subcloning αDB1-VN173 insert from a pBiFC-αDB-VN173 into a pcDNA3.1(+) backbone.

### 2.2. Skeletal muscle preparation and immunostaining

For teased muscle fiber preparation, the fascia covering the *Tibialis anterior* muscle of P250 mice were removed and the legs fixed in 4% PFA (paraformaldehyde) for 40-60 min. The dissected muscle was used to obtain teased bundles of muscle fibers that were permeabilized with a blocking solution (0.2% BSA, 0.5% Triton X-100, 2% normal goat serum) in PBS for 8 h at room temperature. For cryosections, the *Tibialis anterior* muscles from P45 were embedded in OCT, and snap-frozen by direct contact with isopentane cooled in liquid nitrogen and then stored at −80°C until use. Transverse, 20 μm-thick sections were obtained with Sakura CRYO 3 cryostat, collected on super frost slides, dried at room temperature for 1 h, and stored at −20°C until use. For immunohistochemistry, sections were fixed with 4% PFA for 10 min at room temperature and then permeabilized with blocking solution for 1 h.

### 2.3. Antibodies and staining reagents

To visualize muscle cross-sections, Alexa Fluor 555 Phalloidin (# A34055, Thermofisher) and DAPI (# A4099,0005, BUJNO Chemicals) were used. For immunohistochemistry, the following primary antibodies were used: rabbit anti-neurofilament (#AB1991, Millipore); guinea pig anti-synaptophysin1 (#101004, Sysy); rabbit anti-cortactin (#PA5-27134, Invitrogen and #868102, BioLegend). To visualize NMJ compartments, α-bungarotoxin (#B35451, Thermofisher) and Fasciculin-II + ATTO-488 (# F-225, Alomone Labs) were used. Rabbit anti-GFP (# ab32146, Abcam) and rabbit anti-FLAG (DYDDDDK tag) (#14793, Cell Signalling) were used for Western blotting.

### 2.4. Imaging

Immunofluorescence microscopy was performed using a Zeiss Cell Observer SD confocal microscope available at the Bioimaging Laboratory of Łukasiewicz Research Network - PORT Polish Center for Technology Development. The images were collected and analyzed using ZEN 3.1 software (Carl Zeiss) and processed with ImageJ. The automated analysis of NMJs was performed on a z-stack of single NMJ images by NMJ-morph Software [19].

### 2.5. Cell Culture, transfection, and protein extraction

Human embryonic kidney (HEK) 293T cells (# CRL3216, ATCC) were cultured in Dulbecco’s Modified Eagle’s Medium, with 4.5 g/L Glucose, without L-Glutamine, supplemented with 10% fetal bovine serum, 1% Penicillin-Streptomycin (5,000 U/mL) (#15070063, Thermofisher), and 0.1% Amphotericin B (#15290018, Thermofisher). HEK293T cells were transfected at 60-90% confluence with TransIT®-LT1 Transfection Reagent (#2300, Mirus Bio) according to the manufacturer’s protocol. 48h after transfection, the cells were washed with 1x PBS and lysed on ice with the co-IP buffer (50 mM Tris-HCl, 150 mM NaCl, 50 mM NaH2PO4, 10 mM imidazole, 0,1% NP-40, 10% glycerol; pH 8.0) supplemented with Complete Protease Inhibitor cocktail (# 04693116001, Roche). After incubation for 45 min at 4°C with gentle rotation, the homogenates were centrifuged for 20 min at 14.000 rpm at 4°C, and the supernatants collected. C2C12 myoblasts (91031101, Sigma-Aldrich) were cultured as described previously [20]. Briefly, myoblasts were cultured on 0.2% gelatin-coated plates in Dulbecco’s Modified Eagle’s Medium (DMEM) supplemented with 20% fetal bovine serum (FBS), 4.5 g/L glucose, L-glutamine, penicillin, streptomycin, and fungizone. For myotube fusion, cells were plated on petri dishes coated with 111-laminin (L2020-1MG, Merck). To induce myotube differentiation, the media were replaced with DMEM with 2% horse serum. To extract proteins from the myotubes, cells were washed with PBS and scraped from the dish in lysis buffer (50 mM Tris-HCl [pH 8,0], 150 mM NaCl, 0.1% NP-40, 10% glycerol, 1 mM DTT, 1 mM NaF, and supplemented with Complete Protease Inhibitor cocktail. Cell homogenates were pushed five times through 25 gauge needles and incubated at 4°C with gentle rotation for 30 minutes. Next, the lysates were centrifuged at 18 000x g at 4°C for 30 minutes, and the supernatants collected.

### 2.6. Co-immunoprecipitation

Cell lysates were incubated with Dynabeads™ M-270 Epoxy (14301, Thermofisher) pre-coated with mouse anti-GFP (A11120, Life Technology) or mouse anti-Flag (F3165, Sigma) using Dynabeads Antibody Coupling Kit (14311D, Thermofisher) according to the manufacturer’s instructions. Dynabeads were then incubated with the protein extract at 4°C overnight, and washed 3 times with co-IP buffer. Finally, the beads were incubated with 2x Laemmli sample buffer (161-0747, Bio-Rad) for 10 min at 95°C to elute precipitated proteins.

### 2.7. Peptide pulldown and mass-spectrometry

The following synthetic biotinylated peptides were used to immunoprecipitate interacting proteins: aDB1-prep3-730: Biotin-Ahx-RQLENELQLEEYLKQKLQDE-NH2, aDB1-prep3-730-P: Biotin-Ahx-RQLENELQLEEY[11]LKQKLQDE-NH2. Peptides were synthetized at Biomatik as fee for service. Dynabeads® M-280 Streptavidin (65801D, Thermofisher) were incubated with 2µg of each synthetic peptide for 30 minutes at room temperature. After washing with PBS containing 0,1% NP-40 and twice with co-IP buffer, the Dynabeads were incubated with lysates from HEK293T cells transfected with EGFP-cortactin plasmid at 4°C overnight. The beads were washed 3-4 times with co-IP buffer, and the proteins eluted with 2x Laemmli sample buffer (161-0747, Bio-Rad) for 10 min at 95°C. For MS/MS analysis, Epoxy M-270 beads (14301, Invitrogen) were coated with 40 μg of either aDB1-prep3-730 or aDB1-prep3-730-P peptides, and incubated over-night at 4°C with C2C12 homogenates. Next, beads were washed three times with lysis buffer and incubated with 2x Laemmli sample buffer for 10 min at 95°C to elute precipitated proteins. The mass spectrometry analysis was outsourced to the MS Bioworks facility (USA) as fee for service. Briefly, samples were resolved with SDS-PAGE. The electrophoresis lane was divided into ten gel fragments that were used for automated in-gel digestion with a Digilab ProGest robot. Analysis was performed by liquid chromatography coupled with tandem mass spectrometry (LC-MS/MS). Protein identification was performed with Mascot.

### 2.8. Western blot

SDS-PAGE was executed on polyacrylamide gels prepared according to routine protocols. The proteins were transferred onto nitrocellulose membranes (0.2 μm pore size, #1620112, BioRad) using Trans-Blot® Turbo™ Transfer System (Bio-Rad) and blocked in 10% non-fat milk for 1h at room temperature. Primary antibody incubation was performed at 4°C overnight with agitation, while the incubation with secondary antibodies was performed at room temperature for 1h. For detection, we used horseradish peroxidase-conjugated secondary antibodies (Cell Signalling) and Clarity Western ECL substrate (1705060, Bio-Rad) or SuperSignal West Femto Maximum Sensitivity Substrate (34095, Thermo Fischer). ECL images were either developed on photographic X-ray films (771468, Carestream, USA) and digitized using a film scanner (Epson), or acquired by the Gel Doc XR+ System (Biorad).

### 2.9. Animals and genotyping

All of the experimental procedures were approved by the Local Ethics Committee for Animal Experimentation (065/2020/P1 and 066/2020/P1), and the experiments were con-ducted in accordance with all applicable laws and regulations. Cttn-KO mice were as previously published [21]. The animals were maintained at the Animal Facilities at the Łukasiewicz Research Network - PORT Polish Center for Technology Development on 12-hour dark-light cycles with food and water available ad libitum. For genotyping, the genomic DNA was extracted with Chelex 100 Chelating Resin (1421253, BioRad). Briefly, small pieces of biopsies were added to 200µl of 10% Chelex solution, incubated at 951C for 20 min, and centrifuged at 12 000 g at room temperature for 10 min. The supernatant was collected and used as a template in the PCR reactions performed with PCR Master Mix (2X) (K0172, Thermofisher). Primers 5’-AGGGTCTGACCATCATGTCC-3’ and 5’-GTGCTGTTCATCCACAATGC-3’ were used to detect the KO allele, and 5’-CCTGGAATAAGTCAGCCAAGC-3’ and 5’-CGGAGAGCTAGGCTGTTAGC-3’ for the WT allele.

### 2.10. Muscle electroporation

Mice were anesthetized followed by injection of a *Tibialis anterior* muscle with 25 μg of plasmid DNA in 25 μl of PBS using a Hamilton syringe. Electroporation was performed with BTX ECM 830 electroporator (BTX, USA). Electrodes were dipped in PBS before 10 pulses at 150V/cm for 20 ms at 1 Hz were applied. Post-surgery analgesia was achieved by intraperitoneal injection of 0,4 ml/kg of Metacam and topical application of 2% lignocaine gel (lignocainum, Jelfa).

### 2.11. Behavioral tests

To minimize animal stress, mice between three and six months of age were habituated for 5 consecutive days before experiments. The animals’ general locomotor activity was assessed using the Running Wheels (1800/50, Ugo Basile). Each apparatus consists of a cage with a wheel connected to a magnetic counter that records the wheel’s revolutions. The animals were placed during their active period, and left free to move for 12h. Each apparatus was supplied with food and water ad libitum. The pulling strength of mice was tested by the Grip Strength Meter (47200, Ugo Basile), which records the peak-pull force of limbs. Briefly, the animal was allowed to grasp the grid of the Grip Strength Meter and gently pulled backward. The measurement of the maximal force was repeated 10 times per animal with a rest of 60 seconds between each measurement. Endurance to exercise was assessed by a motorized treadmill (57630, Ugo Basile). To minimize stress, animals were allowed to habituate to the device run at 10m/min speed, for 5 minutes, 5 days in a row. The endurance testing started with a 5 minutes warm-up at 10m/min speed, and the treadmill was then automatically and progressively accelerated by 1m/min. Air puffs were employed as an adverse stimulus, which was activated when the animal was standing on the air puff grid instead of running. The exhaustion was considered reached when the animal stopped running and received 15 seconds of consecutive air-puff aversive stimulation.

### 2.12. Statistics

The statistical analysis was performed with GraphPad Prism 6 (GraphPad Software Inc., San Diego, CA, USA) and Microsoft Excel (Microsoft). The results are shown as means ± SEM. The raw data collected were analyzed by unpaired Student’s t-test or one-way ANOVA with Tukey post-hoc correction. Statistical tests that revealed a p-value < 0.05 were considered statistically significant.

## 3. Results

### 3.1. Cttn binding partner of αDB1 at NMJ

Alpha-dystrobrevin-1 (αDB1) has three tyrosines in the C-terminal part (**Figure 1A**) that play important roles in the function of the protein at the neuromuscular junction (NJM) [12]. In a previous study, we have identified several proteins recruited to tyrosine Y713. To identify the molecular interactors of Y730, we used synthetic peptides corresponding to the αDB1 sequence with centrally located Y730. One of the peptide was phosphorylated at the tyrosine residue and the other one was unphosphorylated. Peptides with a biotin residue at the N-terminal end were used to coat magnetic Dynabeads, and to precipitate interacting proteins from C2C12 myotube homogenates. Isolated samples were resolved using SDS-PAGE electrophoresis, trypsin digested and peptides identified by mass spectrometry (**Supplementary Table 1**).

**Figure 1.**
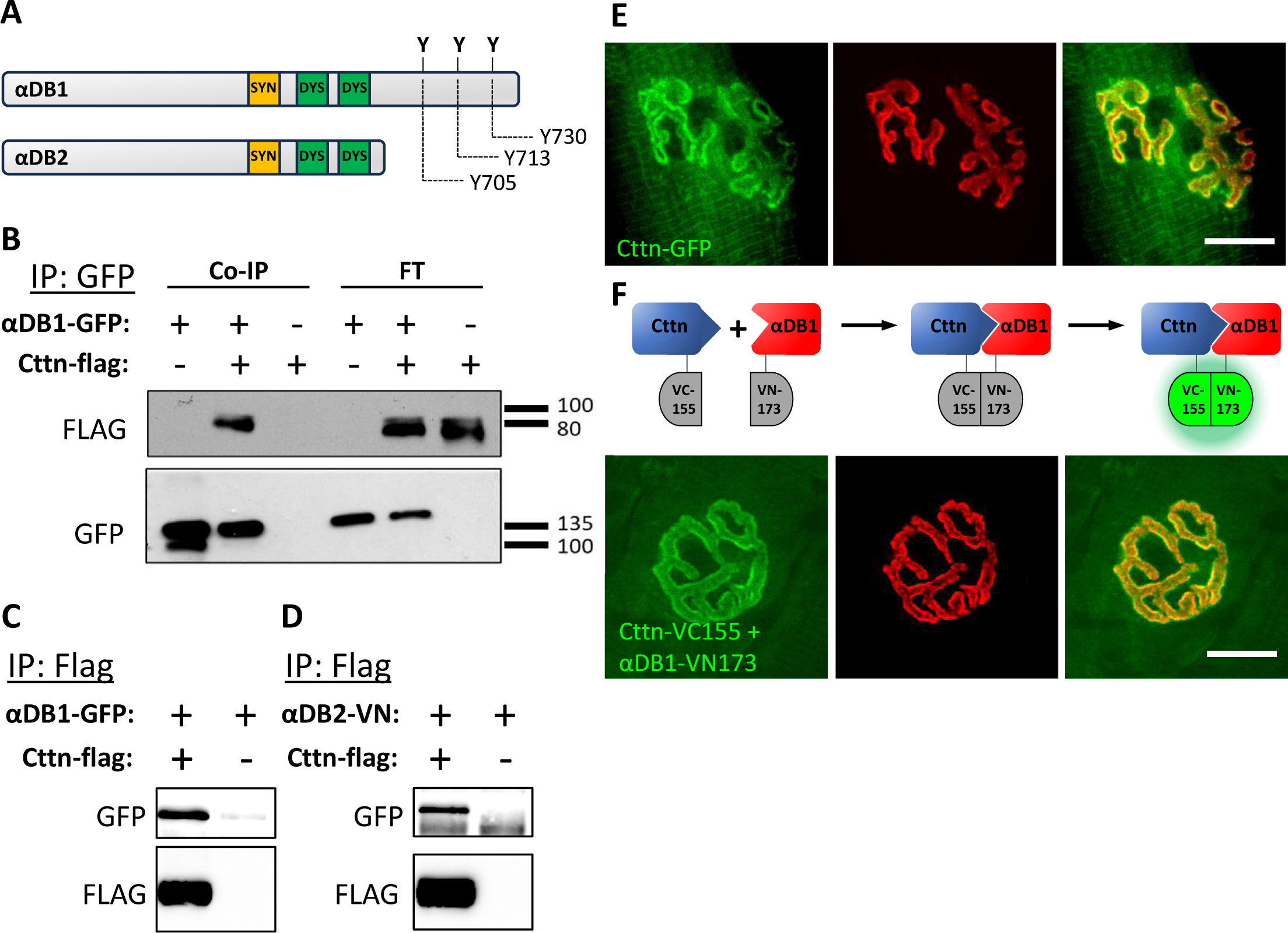
Cttn interacts with αDB1 and localizes to the NMJ postsynaptic machinery. (**A**) Organization of the αDB1 protein domains with highlighted three tyrosines in the C-terminal fragment missing in αDB2. (**B**) αDB1-GFP co-immunoprecipitates Cttn-Flag expressed in HEK293 cells. Cttn-Flag co-immunoprecipitates αDB1-GFP (**C**) and αDB2-VN (**D**) from HEK293 cell homogenat es). **(E)** Cttn-EGFP concentrates at the muscle postsynaptic machinery, co-localizing with the AChR. (**F**) BiFC experiment. The top panel provdies a schematic visualization of the BiFC mechanisms. The lower panel shows the interaction (BiFC signal, green) between Cttn-VC155 and αDB1-VN173. FT - flow through; scale bar 15µm.

This approach revealed 1027 candidate proteins, of which 52 were precipitated specifically with the phosphorylated peptide and 275 with the unphosphorylated one (**Supplementary Table 1**). Among the identified proteins were several important regulators of the actin cytoskeleton (**Table 1**), including two subunits of the Actin Related Protein (Arp2/3) complex, as well as Cyclase Associated Actin Cytoskeleton Regulatory Protein 1 (CAP1), Tyrosine Kinase Substrate with Five SH3 Domains (TKS5), Capping Actin Protein of Muscle Z-Line Subunit Alpha 1 (CAPZA1), Alpha-actinin-2 (ACTN2), and cortactin (Cttn). The Arp 2/3 complex is responsible for the nucleation of branched actin filaments and together with N-WASP, was proposed to regulate the clustering of AChR in response to agrin [22]. The interaction between Tks5 and αDB1 and its potential role in the organization of the AChR had been reported previously [23, 24]. Therefore, in this study we concentrated on the function of Cttn at the NMJ. Cortactin is an F-actin and Arp2/3 complex interacting protein that is implicated in Rho-GTPase signalling and the stabilization of Arp2/3 complex-dependent actin filament branches [25], but it had also been shown to be capable of binding and activating N-WASP [26] Furthermore, cortactin is thought to associate with AChR clusters in cultured muscle cells, where is has been shown to regulate receptor aggregation [27]. However, its localization or function at the NMJ has hitherto remained unclear.

**Table 1.**
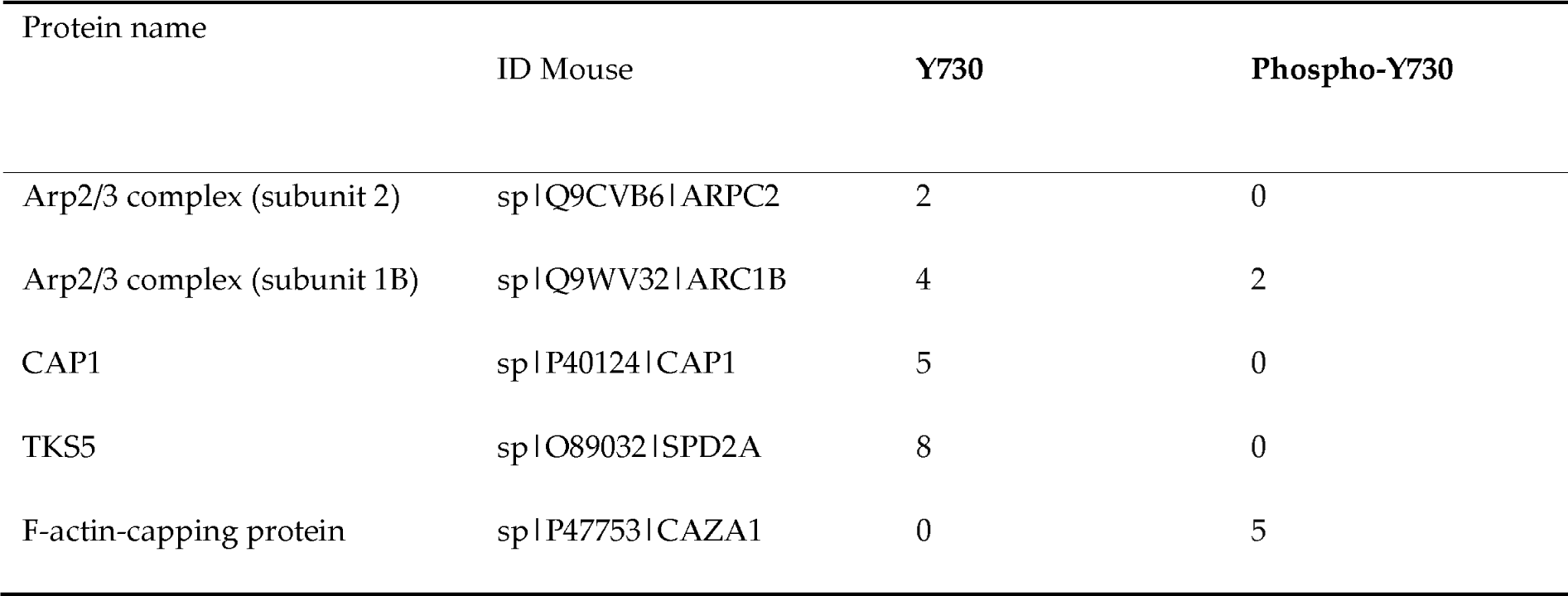
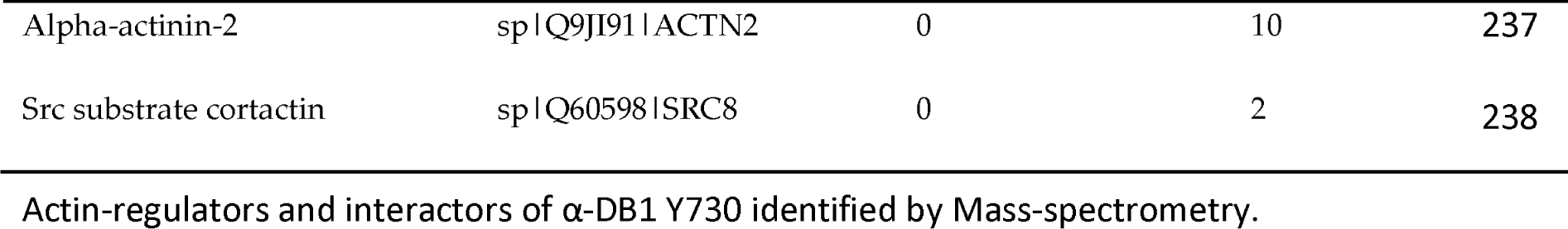
Actin-regulators and interactors of α-DB1 Y730 identified by Mass-spectrometry.

To confirm that αDB1 interacts with Cttn, we performed co-immunoprecipitation experiments of αDB1-GFP (87 kDa) and Cttn-Flag (∼61kDa) expressed in Hek293 cells. As shown in **Figure 1B**, Cttn-Flag was co-precipitated with αDB1-GFP. Similar results were obtained in the reciprocal experiment showing that αDB1-GFP can be co-precipitated with Cttn-Flag (**Figure 1C**), although some residual unspecific binding of αDB1-GFP to the beads coated with the anti-Flag antibody was also detected. These experiments confirmed the interaction between αDB1 and cortactin, which was originally identified in our mass spectrometry experiment. To test if Cttn binds only to the short αDB1 sequence present in the peptide, we decided to co-precipitate Cttn and the αDB2 protein, in which the entire C-terminal fragment of αDB1 with the three tyrosines is missing (**Figure 1A**). In this experiment, the αDB2 was fused to halve of the Venus fluorescent protein that is used in Bi-molecular fluorescent complementation (BifC) assays. This halve of the Venus has amino acid sequence similarities to GFP and can be easily detected by anti-GFP antibody. As shown in **Figure 1C**, we observed the interactions between the two proteins when they are overexpressed in Hek293 cells, suggesting that Cttn can interact with αDB1 in several different protein regions, and not only through its C-terminal part, indicating a more complex interaction mechanism.

### 3.2 Cortactin localizes to the postsynaptic machinery

In a previous study, Cttn was identified to be localized at the site of acetylcholine receptor (AChR) clusters in cultured myotubes [27], but the localization of Cttn in vertebrate NMJ has never been reported. We, therefore, electroporated mouse *Tibialis* muscle with a plasmid encoding EGFP-tagged Cttn. As shown in **Figure 1E**, the EGFP signal was strongly concentrated at the postsynaptic machinery where it completely co-localized with the AChR in the synaptic domains.

To determine the localization of interactions between αDB1 and Cttn, we conducted a Bi-molecular Fluorescence Complementation (BiFC) experiment. For this, we electroporated Cttn-VC155 and αDB1-VN173 into the *Tibialis anterior* muscle. VC155 and VN173 are two fragments of the Venus protein that are non-functional. However, if two interacting proteins bring them in close proximity, the parts of the Venus protein can fold into functional fluorescent protein (**Figure 1F**). Importantly, co-electroporation of split Venus components alone into skeletal muscles does not lead to a BiFC signal at the NMJ [18]. Fluorescent complementation at the junctions is observed only when the two portions of Venus are linked to synaptic proteins that interact with each other bringing VC155 and VN173 in close proximity [18]. As expected, electroporation of Cttn-VC155 and αDB1-VN173 led to a BiFC signal predominantly observed at the NMJ postsynaptic machinery (**Figure 1F**) where Cttn and αDB1 are concentrated.

### 3.3 Cortactin plays an important role in the organization of the NMJ in **Soleus** muscle

It has previously been shown that Cttn plays an important role in coordinating actin-remodelling events that drive the clustering of AChRs in cultured myotubes in response to agrin stimulation and concentrates at the muscle-nerve contact points in the co-culture system [27–29]. To study the function of Cttn at the murine NMJ in vivo, we first analysed neuromuscular synapses in the slow-twitching *Soleus* muscle of systemic Cttn knockout mice. To visualize different synaptic compartments, we stained muscle tissues for postsynaptic AChR, presynaptic synaptophysin present in motor neurons, and the synaptic cleft protein acetylcholine esterase (**Figure 2A**). As shown in **Figure 2A**, the alignment of the presynaptic, postsynaptic, and synaptic cleft proteins in control and Cttn-KO was similar. Furthermore, the morphology of motor neuron axons was normal in the absence of Cttn, as judged by neurofilament (NF) immunoreactivity (**Figure 2C**). Interestingly however, detailed analysis of the synaptic morphology revealed that NMJs in the *Soleus* muscle in Cttn-KO mice are slightly smaller and of abnormal morphology (**Figure 2A**). To quantify this effect, we performed unbiased automated analysis of topological patterns for a large number of NMJs using NMJ-morph plug in to ImageJ [28]. For this, the orthogonal projections of series of confocal optical sections of the NMJs were collected and converted to binary images showing the endplate, the regions occupied by AChRs, and perforations within the NMJs that do not contain receptors (**Figure 2D**). Our analysis confirmed the initial observation that synapse size is smaller on average in Cttn-KO*soleus* muscle compared to control animals of the same age (**Figure 2E**). The decreased junctional size appears to be the consequence of the reduced area occupied by AChR and decreased size of NMJ perforations (**Figure 2F**, **G**). We have also performed analysis of NMJ fragmentation by counting the number of NMJs classified to three categories – class I with fewer than four fragments, class II with four to six NMJ fragments, and class III with more than six fragments (**Figure 2H**). Although a slight trend towards increase of class II and decrease of class III NMJs was observed, analysis of a large number of junctions in WT and Cttn-KO *soleus* muscle has not revealed any statistically significant differences between the two genotypes in respect to NMJ fragmentatio**F**n**ig**(**ure 2H**). Collectively, these results suggest a functional involvement for cortactin in the organization of the neuromuscular synapse in slow-twitching *Soleus* muscle.

**Figure 2.**
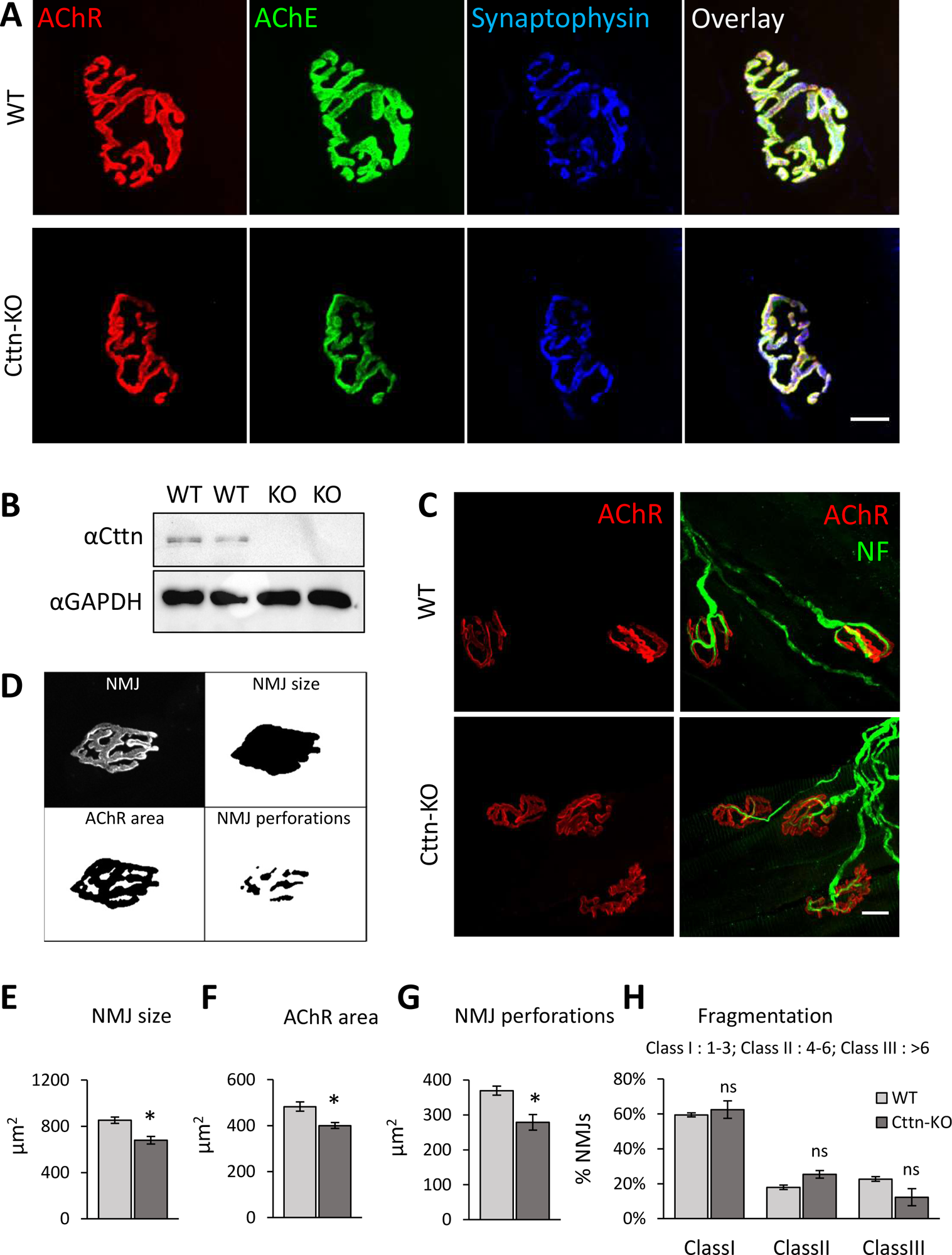
NMJ sizes are decreased in the *Soleus* muscle of Cttn-KO mice. (**A**) Cttn-KO NMJ exhibits normal apposition of the synaptic compartments. (**B**) A Western blot experiment confirming the absence of Cttn protein expression in Cttn-KO *Soleus* muscle homogenate. (**C**) Motor neurons in *Soleus* Cttn-KO have normal morphology, as revealed by neurofilament (NF) immunoreactivity. (**D**) Image processing by NMJ-morph for automated NMJ analysis of size and shape. (**E-G**) Automated quantification of NMJ size (**E**), AChR area (**F**), and NMJ perforations (**G**); unpaired t-test, error bars - SEM. (H) Quantification of NMJ fragmentation in *Soleus* muscles; one-way ANOVA with Tukey post-hoc correction, error bars ‒ SEM; n ≥ 95 NMJs from three animals per genotype. Scale bar = 15µm.

To study the effect of Cttn KO in fast-twitching muscle, we performed a similar set of histochemical experiments on *Tibialis* muscle. Similarly to *Soleus* muscle, we did not detect differences in the alignment of the synaptic components or the morphology of motor neuron processes in *Tibialis* muscle from Cttn-KO and control mice (**Figure 3A**, **B**). Surprisingly, our automated NMJ analysis with the NMJ-morph script revealed that the size of NMJs and the areas of AChR as well as NMJ perforation were only slightly decreased in this case, with the differences not reaching statistical significance (**Figure 3C-E**). Similarly, in spite of a trend towards increase of class I and decrease of class II and III-NMJs, NMJ fragmentation in *Tibialis* muscle of Cttn-KO mice was not statistically significantly different from wild-type animals (**Figure 3F**). We thus conclude that the *Soleus* muscle is more sensitive to the absence of Cttn than the *Tibialis anterior* muscle.

**Figure 3.**
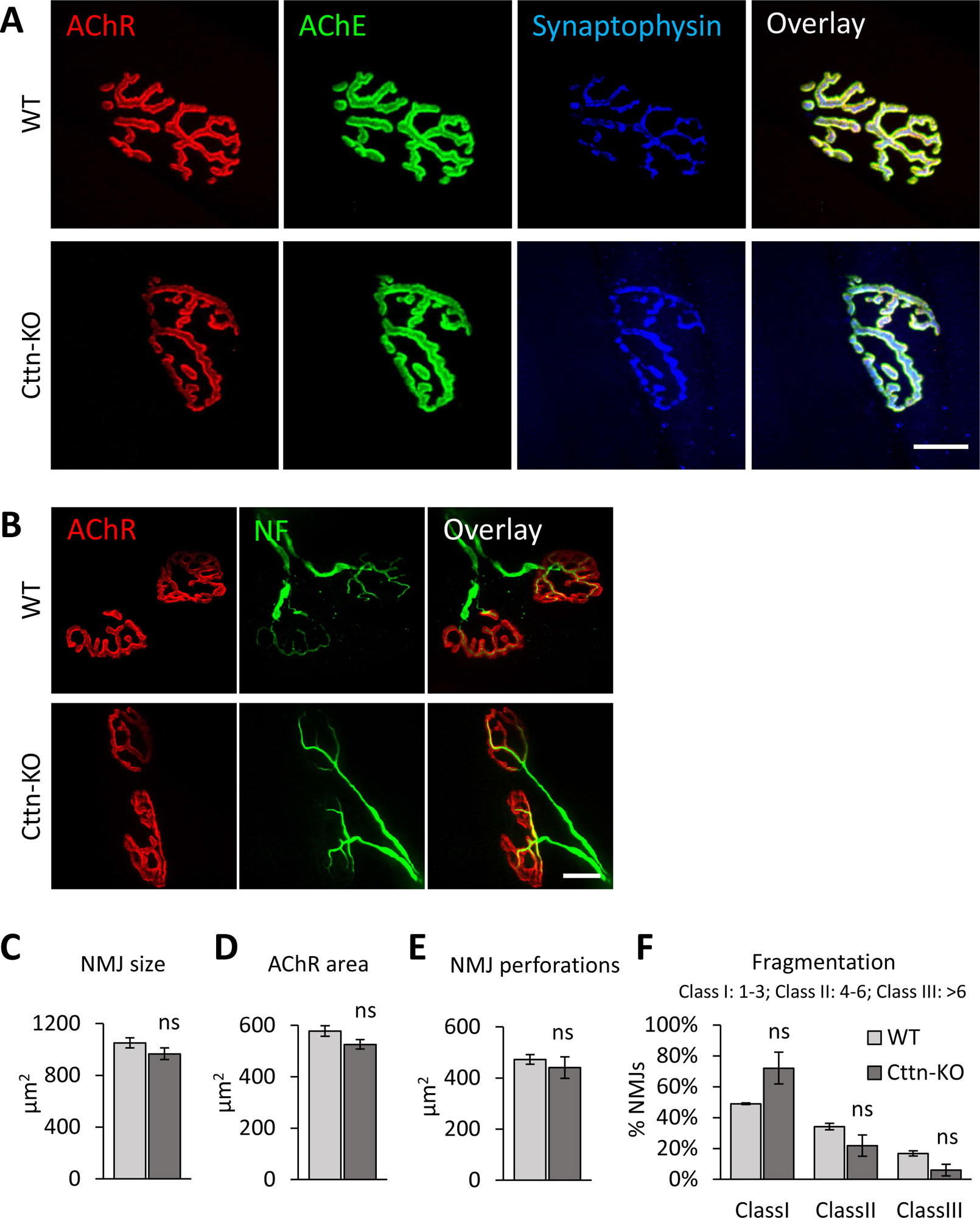
NMJ analysis of*Tibialis* muscle from Cttn-KO mice. (**A**) Cttn-KO NMJ exhibits normal apposition of synaptic compartments; scale bar = 20 μm. (**B**) Normal general morphology of the motor neuron axons in Cttn-KO mice (NF - neurofilament); scale bar = 15μm. (C-E) Automated quantification of NMJ size (**C**), AChR area (**D**), and NMJ perforation area (**E**); unpaired t-test, error bars - SEM. (**F**) Quantification of NMJ fragmentation; one-way ANOVA with Tukey post-hoc correction, error bars ‒ SEM; n ≥ 93 NMJs from three animals per genotype.

### 3.4 Normal muscle fiber size and nuclei localization in Cttn-KO muscles

The DGC components are essential for the integrity of muscle fibers and their dysfunction is associated with the development of dystrophies [7, 14]. Furthermore, morphological abnormalities in the neuromuscular junction (NMJ) may arise from synaptic program dysfunctions or indirectly from muscle fiber atrophy. Consequently, we questioned whether the observed alterations of NMJ structures in the *Soleus* muscle are correlated with muscle fiber degeneration. Therefore, we histochemically analyzed *Tibialis* and *Soleus* muscles cryosections obtained from control and Cttn-KO mice (**Figure 4**). However, microscopic analysis of muscle cryosections with DAPI to visualize nuclei and counter-stained with phalloidin to visualize muscle fibers did not reveal major differences be-tween WT and KO tissues (**Figure 4A**). We than quantified muscle fiber size in *Tibialis* and *Soleus* muscle, but there were again no apparent abnormalities in the absence of Cttn (**Figure 4B**). During fiber degeneration-regeneration cycles, myonuclei migrate from the periphery to the center of the fiber, but in spite of significant experimental variability, in particular for the *Tibialis anterior*, the fraction of fibers with internalized myonuclei was not increased in a statistically significant fashion in both *Tibialis* and *Soleus* Cttn-KO tissues(**Figure 4C**). These results suggest that the lack of Cttn does not have a major impact on the integrity of muscle fibers.

**Figure 4.**
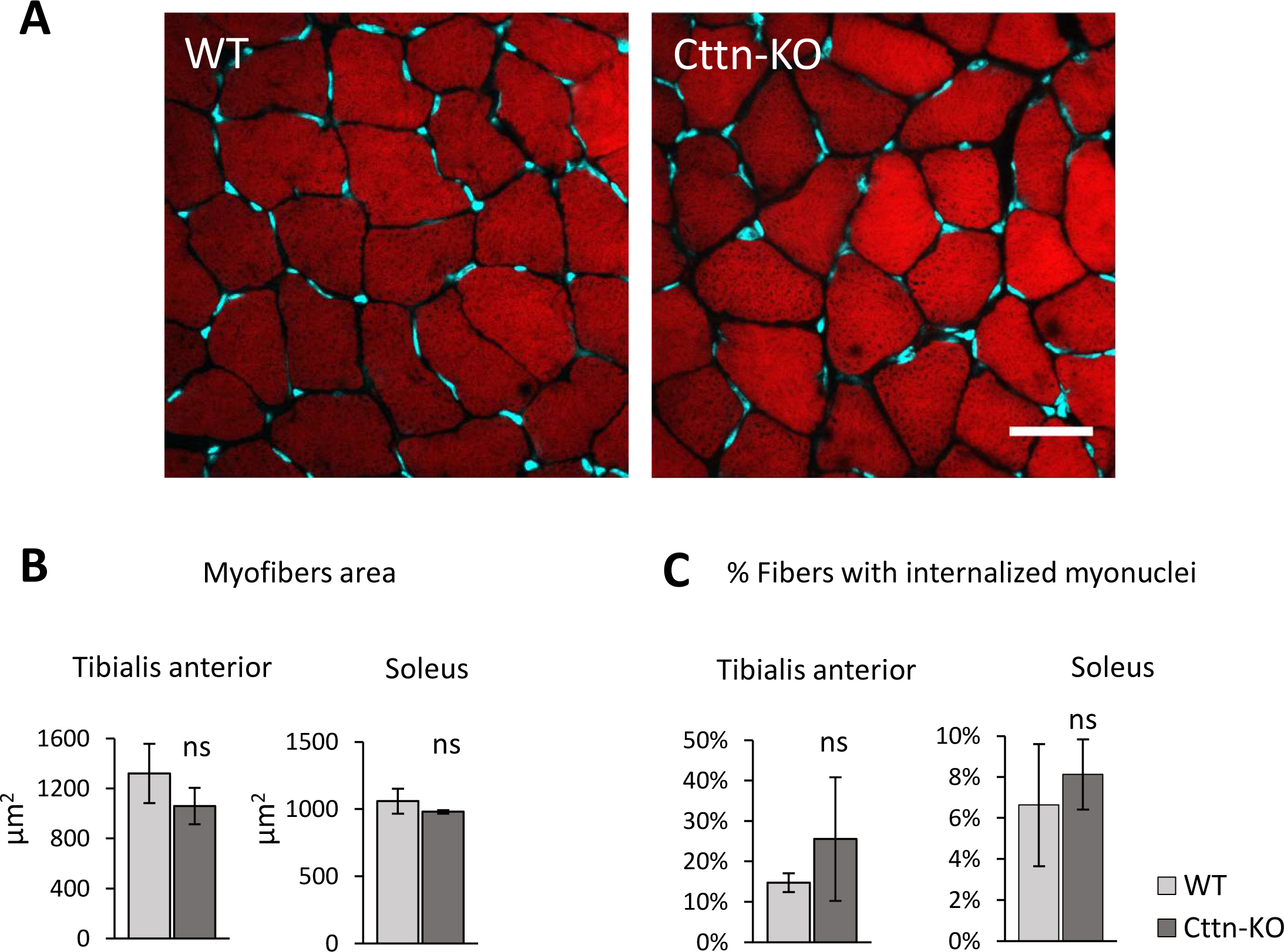
Muscle fiber analysis in the Cttn-KO mice. (**A)** *Soleus* muscle cross-sections, 20x mag, scale bar: 30 µm. (**B**) Fiber size quantification in *Tibialis anterior* (Ta) and *Soleus* (Sol) muscles. (**C**) Quantification of fibers with internalized nuclei in Ta and Sol muscles. Unpaired t-test, error bars ‒ SEM; n ≥ 478 fibers from three animals per genotype.

### 3.5 Loss of Cttn does not lead to a decreased muscle strength or reduced ability of mice to run

To comprehensively investigate the effects of Cttn deficiency on skeletal muscle function, we examined whether the lack of Cttn had any implications on muscle force generation or on the animals’ running ability. Therefore, we conducted a series of tests to assess muscle strength and physical performance of adult Cttn-KO mice (**Figure 5**). The grip strength did not reveal any muscle weakness in Cttn-KO mice (**Figure 5A-C**). To study the ability and willingness of these mice to run, we maintained animals for 12h during the dark phase in a cage with a voluntary running wheel, with the number of rotations of the wheel being recorded at the end of the experiment. Cttn-KO mice ran slightly less indeed on average than WT mice, but the differences did not reach statistical significance. Finally, we monitored the fatigability of these mice when running on an accelerating treadmill, but in this experiment, Cttn-KO mice displayed an identical performance to control animals, with both the distance run until exhaustion and the total time of running being virtually identical in both groups (**Figure 5 E**, **F**). We conclude therefore that the modest alterations observed in NMJ architecture caused by the chronic absence of the actin-binding protein cortactin does not translate into major alterations in muscle strength and physical abilities at least at steady state conditions.

**Figure 5.**
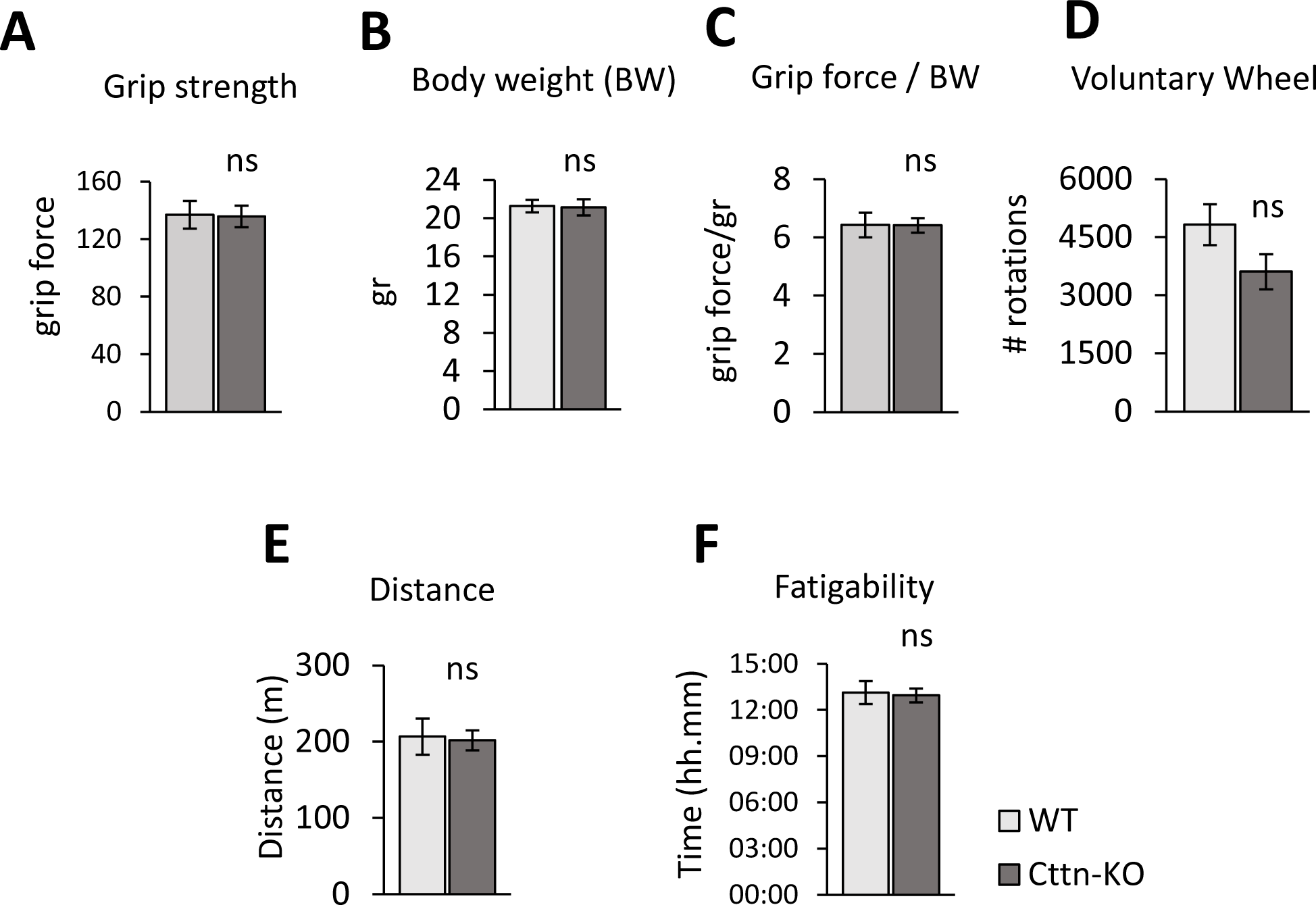
Muscle strength and locomotor abilities of Cttn-KO mice. (**A**) Grip strength test. (**B**) Body weight (BW). (**C**) Grip strength normalized to body weight. (**D**) Number of wheel rotations in voluntary running test. (**E-F**) Treadmill experiment. (**E**) Distance covered over time to reach fatigue. (**F**) Time to reach exhaustion. Unpaired t-test, error bars - SEM, n = 8 mice per genotype.

## 4. Discussion

The preservation of NMJ structural integrity is crucial to ensure efficient synaptic signal transmission, and its disruption has been implicated in the pathogenesis of various neuromuscular diseases [7, 14, 30]. The DGC plays a crucial role in maintaining NMJ integrity and the regulation of the associated cytoskeleton and signalling cascades elicited at neuromuscular junctions. The functions of αDB1, the cytoplasmic component of the DGC that concentrates at the NMJ, are poorly understood, but it was previously reported that the phosphorylation of tyrosine residues at the C-terminal tail of the protein is important [7, 12, 31]. In this study, we report proteomic-based identification of proteins that are recruited to the αDB1 fragment containing Y730 in both modification states, phosphorylated and unphosphorylated. This study identified 1027 candidates, out of which 52 were specifically precipitated with the phosphorylated peptide, and 275 with the unphosphorylated peptide (**Supplementary Table 1**). Several of these proteins likely have important functions at the NMJ. Our initial follow-up experiments focused on the prominent actin cytoskeleton modulator cortactin (Cttn), which can associate with the Arp2/3 complex and the branches it forms during assembly of dendritic actin filament networks. Interestingly, Cttn has previously been demonstrated to play a role in clustering AChR in cultured myotubes in response to agrin stimulation [27]. Cultured Cttn-depleted cells made fewer AChR clusters as compared to controls. This agrees with several other studies demonstrating that the clustering of AChRs relies on the proper remodelling of the actin cytoskeleton. Nevertheless, the localization and function of Cttn at the NMJ in vivo have not been explored. Our studies use biochemical approaches demonstrating that Cttn indeed interacts with αDB1. We show that Cttn is localizing to the murine NMJ, where it concentrates at the postsynaptic machinery, the place likely of tight association with αDB1. Collectively, our studies highlight the function of αDB1 and its phosphorylation as a hub for recruiting signalling and cytoskeleton organizing proteins including Arhgef5, Tks5, Grb2, Liprin, Catulin, and Cttn, all of which are proposed to play important functions at the NMJ [8, 15, 23, 32, 33].

We find here that systemic deletion of Cttn leads to mild alterations in NMJ morphology of the *Soleus* muscle, including the junction’s abnormal shape, decreased endplate size and reduced areas occupied by AChR. Similar tendencies were observed in the *Tibialis* muscle, but in this case, the differences in average values of endplate size and AChR area between WT and Cttn-KO did not reach statistical significance. The *Soleus* and *Tibialis* muscles are composed of two distinct classes of fiber types. The *Soleus* comprises predominantly slow-twitching fibers, while the *Tibialis* contains mostly fast-twitching fibers. They express different Myosin heavy chain (MHC) isoforms. Myofibers of isoform type I (MHCI) exhibit slow contraction speed, while muscles of isoform type II (MHCIIA, MHCIIX, MHCIIB) are classified as fast-contracting. The different contractile properties display differential metabolic requirements; thus, fast fibers of *Tibialis* are efficiently fueled by glycolytic metabolism, while slow fibers rely on oxidative metabolism [34]. Due to variations in fiber composition, distinct skeletal muscles serve differential purposes. The *Soleus* muscle functions as a movement stabilizer, which possesses lower contractile power, but exhibits superior endurance to fatigue. On the other hand, the fast-glycolytic fibers of the *Tibialis* muscle are rapidly recruited during locomotion, but tend to fatigue more quickly.

Similarly, NMJs in the fast and slow muscles can exhibit differences in their general morphology and size [35]. At the molecular level, NMJs in both muscle types appear similar. Still, some differences were reported showing that the synaptic vesicle protein (SV2A) is selectively expressed at the slow-motor axon terminals in adult mice [36]. It has also been reported that some proteins exert differential functions in the different muscle types: for example, the deficiency of the Vesicular ACh Transporter (VAChT), involved in cholinergic transmission, can have different impact on slow- and fast-twitching muscles [37]. Previous studies, therefore, could explain why Cttn deficiency had a differential effect on the *Soleus* and *Tibialis* muscles. The underlying mechanism could involve more efficient compensatory mechanisms to the absence of cortactin in fast muscles or rely less perhaps on the precise regulation of actin cytoskeleton dynamics. NMJ is a highly adaptable structure that has developed compensatory mechanisms to uphold functionality and integrity of the neuromuscular system, which is vital for survival. It is also possible that other molecules could compensate for the lack of Cttn functions. Of note, cortactin is considered a class II nucleation promoting factor and binding F-actin instead of G-actin, the latter of which is essential for the highly efficient Arp2/3 complex activation by canonical, class I nucleation promoting factors, such as N-WASP [38, 39]. So cortactin may have more specific, modulatory functions in Arp2/3-dependent actin assembly, which would also be consistent with its non-essential role in lamellipodium protrusion and actin assembly processes accompanying endocytosis [21, 25, 40].

## 5. Conclusions

Our studies identified cortactin as a novel binding partner of αDB1, the cytoplasmic component of the DGC. We show that cortactin is enriched at the NMJ postsynaptic machinery and can be essential in regulating junctional morphology. Although the NMJ phenotype in Cttn-KO mice is relatively mild, and we observed no decrease in muscle strength in mutant animals, it is possible that Cttn could be indispensable in various pathological conditions, contributing substantially to the maintenance of the NMJ and muscle integrity.

Further research should ascertain the extent of the potential roles of cortactin itself, or compensatory mechanisms following its depletion, and their significance under conditions of NMJ injury, in NMJ autoimmune diseases, or congenital myasthenic syndrome.

## List of abbreviations

ACh: acetylcholine
AChR: acetylcholine receptor
ACTN2: alpha-actinin-2
Arhgef5: Rho Guanine Nucleotide Exchange Factor 5
Arp2/3: actin related protein 2/3 complex
BiFC: bimolecular fluorescence complementation
BSA: bovine serum albumin
BTX: bungarotoxin
Cap1: cyclase-associated protein 1
CAPZA1: capping actin protein of muscle Z-line subunit alpha 1
Cttn: cortactin
Cttn-KO: cortactin knock-out
DAPI: 4’,6-diamidino-2-phenylindole
DGC: dystrophin-associated glycoprotein complex
DMEM: Dulbecco’s Modified Eagle’s Medium
DNA: deoxyribonucleic acid
ECL: enhanced chemiluminescence
ECM: extracellular matrix
EGFP: enhanced green fluorescent protein
FBS: fetal bovine serum
GFP: green fluorescent protein
GRB2: growth factor receptor-bound protein 2
HEK: human embryonic kidney 293 cells
IP: immunoprecipitation
LC-MS/MS: liquid chromatography coupled with tandem mass spectrometry
Lrp4: LDL receptor-related protein 4
MHC: myosin heavy chain
MS/MS: tandem mass spectrometry
MuSK: muscle-specific kinase
NF: neurofilament
NMJ: neuromuscular junction
nNOS: neuronal nitric oxide synthase
NP-40: octylphenoxypolyethoxyethanol
N-WASP: neural Wiskott-Aldrich Syndrome protein
OCT: medium for optimal cutting of frozen tissue
PBS: phosphate buffer saline
PFA: paraformaldehyde
SDS-PAGE: sodium dodecyl-sulfate polyacrylamide gel electrophoresis
SV2A: synaptic vesicle protein 2A
TKS5: tyrosine kinase substrate with five SH3 domains
VAChT: vesicular ACh transporter
αDB: alpha dystrobrevin

## Declarations

### Institutional Review Board Statement

The animal study protocol was approved by the Local Ethical Committee for Animal Experimentation (permission number 065/2020/P1 and 066/2020/P1).

### Consent for publication

Not applicable.

### Data Availability Statement

Data are available from the corresponding author upon request.

### Author Contributions

Conceptualization, T.D.C., M.P., O.W., and T.J.P.; methodology, T.D.C., M.P., O.W., K.R., and T.J.P.; investigation, T.D.C., M.P., O.W., K.R., and T.J.P.; resources, T.D.C., M.P., O.W., K.R., and T.J.P.; writing—original draft preparation, T.D.C. and T.J.P.; writing—review and editing, T.D.C., O.W., K.R., and T.J.P.; funding acquisition, T.J.P. and T.D.C.. All authors have read and agreed to the published version of the manuscript.

## Acknowledgements

Not applicable.

---

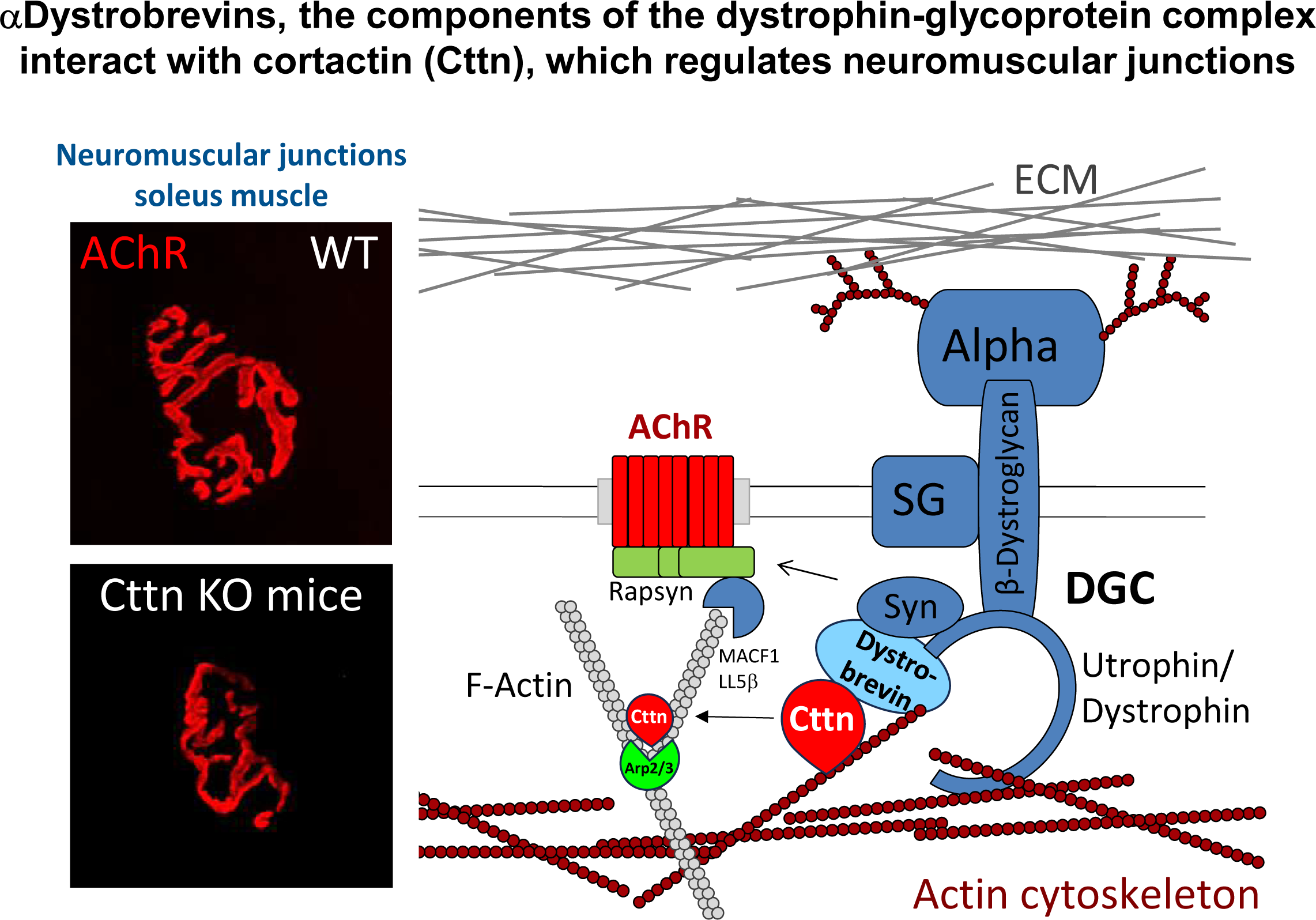

